# Elucidation of the determinant for orchestration of solo unisexual cycle in an important human fungal pathogen

**DOI:** 10.1101/867408

**Authors:** Pengjie Hu, Huimin Liu, Lei Chen, Guang-Jun He, Xiuyun Tian, Xiaoxia Yao, Tong Zhao, Chunli Li, Changyu Tao, Ence Yang, Linqi Wang

## Abstract

In fungi, the sex-determination program universally directs sexual development and syngamy (the fusion of gametes) that underlies pre-meiotic diploidization. However, the contribution of sex-determination to syngamy-independent sexual cycle, which requires autopolyploidization as an alternative approach to elevate ploidy before meiosis, remains unclear in fungi and other eukaryotes. The human fungal pathogen *Cryptococcus neoformans*, as a model organism for studying fungal sexual reproduction, can undergo syngamy-dependent bisexual and syngamy-independent solo unisexual reproduction, in which endoreplication is considered to enable pre-meiotic self-diploidization. Here, by characterizing a mutant lacking all the core sex-determination factors, we show that sex-determination plays a central role in bisexual syngamy but is not strictly required for unisexual development and self-diploidization. This implies an unknown circuit, rather than the sex-determination program, for specifically coordinating *Cryptococcus* unisexual cycle. We reveal that syngamy and self-diploidization are both governed by the Qsp1-directed paracrine system via two regulatory branches, Vea2 and Cqs2. Vea2 directs bisexual syngamy through the sex-determination program; conversely, Cqs2 is dispensable for bisexual syngamy but activates unisexual endoreplication. Through functional profiling of 41 transcription factors documented to regulate *Cryptococcus* sexual development, we reveal that only Cqs2 can drive and integrate all unisexual phases and ensure the production of meiospore progenies. Furthermore, ChIP-seq analysis together with genetic evaluation indicate that Cqs2 induces unisexual self-diploidization through its direct control of *PUM1*, whose expression is sufficient to drive autopolyploidization. Therefore, Cqs2 serves as the critical determinant that orchestrates *Cryptococcus* multistage unisexual cycle that does not strictly require the sexual-determination program.

## Introduction

A sex-determination system is used for the development of sexual characteristics in evolutionally diverse eukaryotes, including fungi [1]. In fungi, the sex-determination system universally directs syngamy (the fusion of gametes to enable pre-meiotic diploidization) and sexual differentiation events during either heterothallic or homothallic mating [2, 3]. However, syngamy does not always underlie sexual development and the two processes can be unconnected in many eukaryotes [4]; the contribution of sex-determination to the fitness related to syngamy-independent solo sexual cycle has yet to be investigated in fungi or other eukaryotes. *Cryptococcus neoformans* is a model organism for studying fungal sexual reproduction. This fungus is a prevalent human fungal pathogen causing more than 200,000 new human infections annually [5], and sex is hypothesized to contribute to its infections through meiosis-created lineage advantages [6–10] and infectious meiospores [11–17]. *C. neoformans* has two mating types (α and **a**) and can undergo two sexual cycles, α-**a** bisexual and unisexual reproduction (also named haploid fruiting), which involves only a single mating type (mainly α) [18–22]. Due to an extreme bias of α in populations (>99%) in nature that may result in infrequent bisexual mating, α unisexual reproduction is considered to be the major sexual form in *C. neoformans* [22–24]. Cryptococcal unisexual reproduction can be differentiated from bisexual reproduction by molecular and morphological events [21, 25]. For bisexual reproduction, the **a** and α cells recognize each other through the pheromone paracrine system and undergo syngamy, forming heterokaryons that subsequently develop into dikaryotic hyphae and allow pre-meiotic diploidization (Figure 1A) [21, 25]. In contrast, syngamy barely happens during unisex [25]. Alternatively, an unusual autopolyploidization process, named endoreplication, is presumed to be the major approach that enables diploidization prior to meiosis during the syngamy-independent unisexual cycle (Figure 1A) [25]. However, the key regulatory basis underlying unisex-specific self-diploidization remains poorly understood.

**Figure 1.**
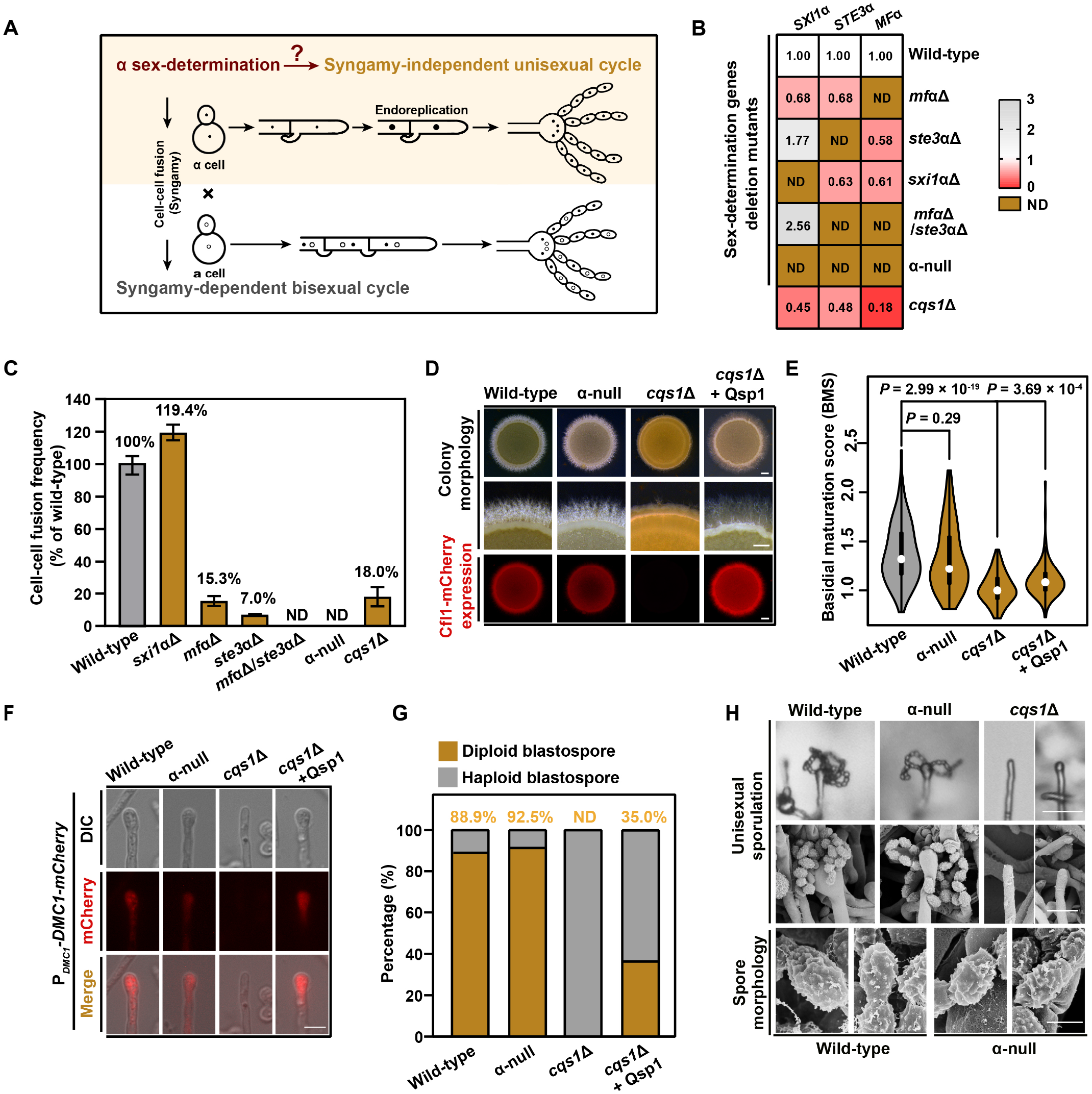
Sex-determination is largely dispensable for unisexual reproduction in *C. neoformans*. **(A)** Schematic diagram depicting unisexual and bisexual development in *C. neoformans*. Endoreplication and syngamy represent two major approaches underlying pre-meiotic diploidization during unisexual and bisexual reproduction, respectively. **(B)** The expression of the α-determining genes in different mutants at 12 hours post unisexual stimulation. Data are presented as the mean of six independent experiments. The color bar represents fold change. ND, not detected. **(C)** Cell-cell fusion frequency of different mutants compared to wild-type. Bilateral cell-cell fusion assay was performed in the *cqs1*Δ mutant, and unilateral cell-cell fusion assay was carried out in other mutants. Data are presented as the mean ± SD of three independent experiments. **(D)** The expression of Cfl-mCherry in various mutants during unisexual community development. The strains were incubated on V8 agar for 7 days at 25°C in the dark. Scale bar, 1mm. **(E)** Violin plot analysis shows that disruption of *CQS1* but not the α-determining genes led to a decrease of high BMS basidial population during unisexual development. Hyphae with or without basidia were photographed at 7 days after unisexual stimulation, and were randomly chosen for the BMS calculation (*N* = 150 for each strain). Two-tailed Student’s *t*-test. **(F)** The expression of Dmc1-mCherry in basidia of α-null and *cqs1*Δ compared to wild-type. The images of the fluorescence-labeled strains were taken at 7 days post inoculation on V8 agar. Scale bar, 10 μm. **(G)** FACS-based ploidy estimation of blastospores dissected from unisexual hyphae of different *Cryptococcus* strains. For each strain, 40 blastospores dissected from different unisexual hyphae were tested for ploidy level. **(H)** Unisexual sporulation phenotypes for wild-type, α-null and *cqs1*Δ strains. Scale bars, 10 μm (upper panel), 5 μm (middle panel), 1 μm (bottom panel).

Despite a remarkable difference between two reproduction modes with regard to their requirement of syngamy, they share multiple development and molecular processes, and are initiated in response to the same environmental cues, which can be achieved through transferring unisexual cells (unisex) or premixed opposite mating cells (bisex) onto sex-inducing medium [21, 23]. After sexual induction, the morphological transition from yeast to hyphae occurs in a stochastic manner [26]. A subpopulation of aerial hyphae subsequently differentiates to form sexual structure (basidia) at their tips. Basidial differentiation occurs concomitantly with meiotic progression, ultimately leading to the production of four spore-chains from the basidium apex [16, 21]. These processes in unisexual and bisexual reproduction are controlled by gene networks with similar architectures. In this regard, the mating mitogen-activated protein kinase (MAPK) pathway is believed to be the core cascade [14, 23]. In this pathway, the downstream MAPK kinase cascade components direct cellular responses to sexual cues through the mating output regulator Mat2 during both unisexual and bisexual reproduction [14, 27, 28]. In contrast to these elements that are essential for both sexual forms, some mating MAPK components appear to have different contributions to the two reproduction modes. For example, the regulatory specificity of bisexual events including syngamy or dikaryotic filamentation is associated with the mating MAPK pathway genes that encode the pheromone Mfα, the Ste3-type pheromone receptor Ste3α and the homeobox regulator Sxi1α [29]. These three factors have been demonstrated to be sufficient to specify the α cell type in *C. neoformans* and represent the key sex-determination components [29]. Notably, independent studies have indicated that the absence of these sex-determination factors individually does not markedly affect unisexual development, in contrast to their important role during the bisexual cycle [30–33]. These findings raise two possibilities regarding the contribution of sex-determination to syngamy-independent unisex that remain to be addressed: i) there is a functional redundancy among these α identity factors during unisexual reproduction, which requires the sex-determination determined by their collective action; ii) the sex-determination established by these factors is not essential for *Cryptococcus* unisexual cycle, which requires an alternative determinant for its initiation and coordination.

## Results

### Sex-determination is largely dispensable for unisexual development in *C. neoformans*

To address the above-mentioned possibilities, we utilized the wild-type strain XL280α, the model isolate for studying *Cryptococcus* unisexual reproduction [19, 34], to generate a mutant (referred to as α-null mutant) that lacks all five genes previously documented to encode the essential α sex-determination factors (Figures 1A and 1B) [29]. These genes include *STE3*α and *SXI1*α as well as three paralogue genes encoding the pheromone Mfα. Compared to each of the single α identity factor mutants, the α-null mutant displayed a complete defect of **a**-α cell fusion during bisexual reproduction (Figure 1C), thereby demonstrating the pivotal role of sex-determination in dictating bisexual syngamy in *C. neoformans*. We next sought to evaluate the role of α sex-determination during unisexual development. To this end, the α-null mutant was quantitatively or semi-quantitatively examined for its ability to undergo sequential unisexual differentiation processes (Figures S1, and 1D-1H). Our results demonstrated that the removal of α identity genes did not markedly impair unisexual filamentation and the expression level of mCherry-tagged Cfl1 in unisexual communities, which is an indicator protein for filamentous development reported previously (Figure 1D) [35]. Similarly, in the α-null mutant, no detectable defect was identified during basidial maturation, based on the basidial maturation score (BMS) assay that allow us to quantitatively evaluate basidial maturation (Figure 1E) [16]. In addition, we showed that the α sex-determination system did not appear to impact meiosis during unisexual reproduction, since the simultaneous deletion of the α sex-determination genes cannot cause evident changes in the spatiotemporal expression of Dmc1 (a meiosis-specific recombinase), a meiosis indicator protein that is highly abundant during meiotic progression (Figure 1F) [36]. We further tested whether the α-identity genes are important for unisex-specific endoreplication using an approach reported previously [25], in which blastospores produced along hyphae from different strains were applied to a FACS-based assay for the assessment of their ploidy status. We found that 92.5% blastospores from the α-null mutant were diploid and this ratio is comparable to that of the wild-type strain, indicating that sex determination is not required for unisex-specific endoreplication (Figure 1G). Significantly, the formation of four spore chains was readily detected when culturing the α-null mutant itself on V8 agar to induce a unisexual cycle (Figure 1H). Additionally, scanning electron microscopy (SEM) experiments did not show any detectable morphological difference between the unisexual spores from wild-type and the α-null mutant (Figure 1H). These lines of evidences indicate that the α unisexual cycle can be successfully and efficiently initiated and accomplished in *Cryptococcus* strains without α sexual identity.

### Qsp1 is the central intercellular signal for unisexual self-diploidization

Our mating efficiency assay indicated that α pheromone and Ste3α represent the core constituents of the α sex-determination system during bisexual syngamy, since simultaneous deletion of their coding genes was sufficient to completely block α-a cell fusion (Figure 1C). In contrast, unisexual development was largely unaffected in α null mutant, in which both Mfα and Ste3α are absent (Figures 1D-H). Unlike these components of pheromone paracrine system that are not essential for unisex, we have previously established that the intercellular cascade directed by Qsp1, a quorum sensing peptide [37, 38], acts as the important intercellular pathway that affects unisexual filamentation and sporulation via an atypical zinc-finger regulator Cqs2 [39]. BMS analysis also indicated that the absence of Qsp1 greatly reduced the population of mature basidia during unisex (Figure 1E), suggesting its engagement in basidial maturation. To further corroborate the importance of Qsp1 in unisexual reproduction, we examined the impact of Qsp1 on unisex-specific endoreplication (Figure 1G). Surprisingly, we failed to identify diploid blastospores dissected from up to 40 unisexual hyphae, in which Qsp1 is absent. Moreover, the synthesized Qsp1 peptide can partially rescue the defective autopolyploidization, supporting that Qsp1 plays a predominant role in the paracrine control of self-diploidization during unisexual reproduction.

### Qsp1 is not specific for the unisexual cycle and evokes α-a bisexual syngamy through the Velvet-family regulator Vea2

Intriguingly, deletion of the Qsp1 coding gene *CQS1* also resulted in a significant decrease in **a**-α cell fusion efficiency, and this defect can be partially restored by the addition of synthetic Qsp1 molecule (Figure 2A). This result indicates that Qsp1 also controls bisexual syngamy. This conclusion is consistent with the transcriptional evidence showing drastically reduced expression of all core sex-determination genes in the absence of Qsp1 (Figures 1B and S2A). Unexpectedly, the **a**-α cell fusion efficiency was not markedly altered in the absence of Cqs2, as revealed by **a**-α bilateral mating analysis (Figure 2A), suggesting that unlike Qsp1, Cqs2 is not important for bisexual syngamy. After reanalyzing the RNA-seq data reported previously [39], we found that deletion of Cqs2 cannot significantly change the mRNA levels of the α sex-determination genes (Figure S2A). These findings indicate a considerable dissimilarity of the stimulatory effects of Qsp1 and Cqs2 on bisexual syngamy and the expression of bisex-specific sex-determining genes, implicating that additional Qsp1-responsive transcription factors (TFs) responsible for these processes likely exist. If so, then such TFs should be controlled by Qsp1 but not Cqs2. In comparing available RNA-seq data derived from Qsp1 depletion versus Qsp1 repletion condition [39], we identified 11 TF coding genes whose expression is considerably upregulated by Qsp1 (Table S1). Among these genes, the expression of 7 TF genes was not significantly changed in the absence of Cqs2. (Table S1). These genes include *SXI1*α, which has been shown to be not essential for bisexual syngamy [40], and the other 6 genes were mutated individually in the congenic XL280α and XL280**a** strains for the assessment of their impact on mating efficiency via bilateral mating evaluation. We showed that 4 out of the 6 TF genes tested dramatically affected α-**a** cell fusion efficiency (Figure 2B). Among these 4 TFs, only absence of Mat2 and the Velvet family regulator Vea2 completely abolished bisexual syngamy (Figure 2B) and significantly reduced the expression of the sex-determination genes, based on quantitative real-time PCR (qRT-PCR)-guided transcription assays (Figure 2C). Vea2 was identified previously by *Agrobacterium tumefaciens*-mediated insertional mutagenesis analysis that showed a defective filamentation caused by a T-DNA insertion in the *VEA2* gene [33], but its role in bisexual syngamy was unknown. We found that the addition of excessive synthetic Qsp1 (512 nM) cannot restore the cell fusion defect in the *vea2*Δ mutant, indicating its role as a key regulatory branch downstream of Qsp1 during bisexual syngamy (Figure S2B). Given that both Vea2 and Mat2 are required for bisexual syngamy and the sex-determination gene expression and filamentous growth, we proposed a potential regulatory relationship between these two regulators. As mentioned above, Mat2 is considered to direct syngamy as the terminal component of *Cryptococcus* mating signaling cascade [28]. Therefore, it is conceivable that Vea2 may function upstream of Mat2 during bisexual syngamy. This idea was verified through the transcriptional evidence revealed by qRT-PCR assays that Vea2 significantly upregulated the expression of Mat2 but not *vice versa* (Figure S2C).

**Figure 2.**
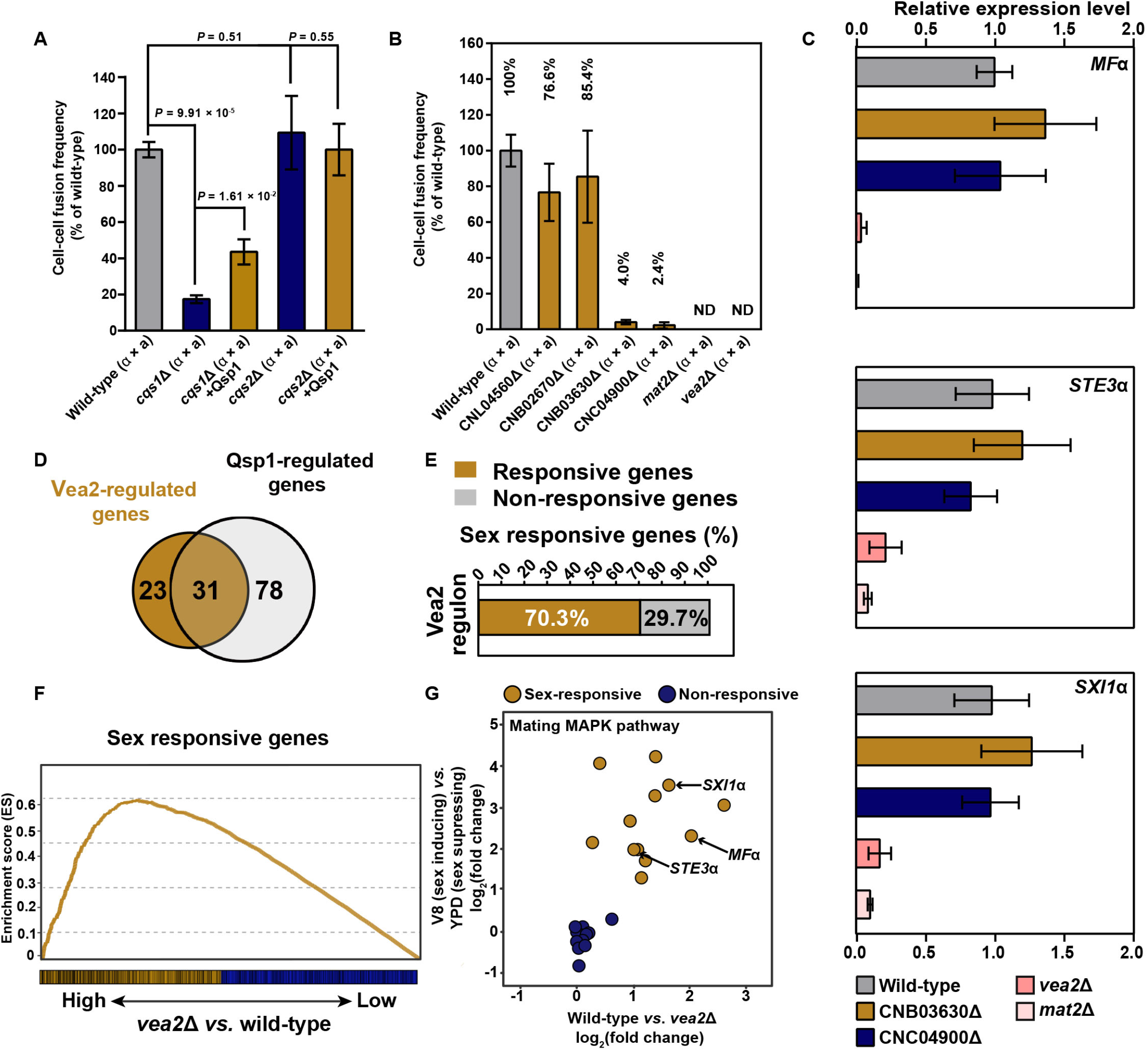
Qsp1 controls α-a bisexual syngamy through the velvet family regulator Vea2. **(A)** Bilateral *cqs1*Δ and *cqs2*Δ mutant cell fusion frequency in the presence and absence of synthetic Qsp1peptide (512 nM) compared to wild-type. *N* = 3 independent experiments, mean ± SD, two-tailed Student’s *t*-test. **(B)** Bilateral cell fusion frequency of the Qsp1-responsive TF mutant strains. Data are presented as the mean ± SD of six independent experiments. TF, transcription factor. ND, not detected. **(C)** The relative mRNA levels of the sex-determining genes in the Qsp1-responsive TF mutant strains at 12 hours post inoculation on V8 medium. *N* = 6 independent experiments, mean ± SD. **(D)** Venn diagram analysis indicating a large overlap between the targets regulated by Qsp1 and Vea2. **(E)** Vea2 controls many sex-responsive genes. **(F)** Gene-set enrichment analysis (GSEA) showing a significant enrichment of sex-responsive genes in the Vea2 regulon. Vertical lines on the X axis are members of the gene set, ordered by value of log_2_ (*vea2*Δ *vs*. wild-type). **(G)** Many mating MAPK genes induced by sex stimuli are activated by Vea2. The arrow indicates the α sex-determining genes.

To identify the regulon of Vea2, we utilized a high-coverage strand-specific RNA-sequencing analysis to compare whole-genome expression between wild-type and the *vea2*Δ mutant at 12 hours after sexual induction, when a majority of genes responsible for early sexual events (such as syngamy) are induced based on our previous study [16]. We identified 54 protein-coding genes that displayed markedly altered expression in the absence of Vea2 (Table S2, log_2_|fold-change| > 1.0, q value < 0.01, TPM (transcripts per million mapped) > 5). Significantly, there is a great overlap between the targets controlled by Qsp1 and Vea2, respectively (*P* = 5.44 × 10^−4^, Fisher’s exact test), verifying the role of Vea2 as an important target of Qsp1 (Figure 2D). We found that ~70.3% of the Vea2 regulon exhibited evident transcriptional response to the external cue that induces cryptococcal sex (*P* = 3.10 × 10^−4^, χ2), pointing to the specificity of Vea2 in cellular response to sex-inducing cues (Figure 2E). This conclusion is further corroborated by GSEA analysis, which showed a significant enrichment of early sex-responsive genes among the Vea2 regulon (*P* < 0.01, permutation test) (Figure 2F).

Especially, many genes induced by Vea2 belong to the mating MAPK pathway, the key cascade for early mating response and syngamy, whereas none of Vea2 targets are from other well-known cascades (Figure S2D). Furthermore, the expression of all α identity genes were significantly reduced in the absence of Vea2 (Figure 2G), echoing the qRT-PCR results (Figure 2C). These data together support the requirement of Vea2 as a downstream circuit of the Qsp1 cascade for the expression of sex-determination genes and bisexual syngamy.

We next examined whether Vea2 is also critical for unisexual development. Despite attenuated self-filamentation as shown previously [33], the deletion of *VEA2* does not appear to affect late events of the unisexual cycle. Quantitative fluorescence imaging revealed no remarkable change in the expression of mCherry-tagged Dmc1 within basidia during unisex in the absence of Vea2 (Figure S2E). Furthermore, post-meiotic sporulation can be detected when the α *vea2*Δ mutant was grown alone on V8 agar to stimulate the unisexual cycle (Figure S2F).

### Cqs2 is the regulatory determinant underlying unisexual development and self-diploidization

The dispensability of Cqs2 in syngamy, a process specifically essential for bisexual reproduction, encouraged us to test the importance and specificity of Cqs2 during the unisexual cycle. The influence of Cqs2 on endoreplication was first examined because this process is specific for syngamy-independent unisexual reproduction. FACS analysis revealed that 40 blastospores produced from the *cqs2*Δ mutant were all haploid (Figure 3A). This indicates that Cqs2 is crucial for *C. neoformans* to undergo unisexual endoreplication. The essential function of Cqs2 on unisexual self-diploidization suggests the possibility that Cqs2 may serve as the master regulator of the unisexual cycle. To address this possibility, we exploited the Cqs2 overexpression system (the P_*H3*_-*CQS2* strain) to determine whether the intact multistage unisexual cycle can be properly promoted and coordinated in response to its increased expression. To our expect, Cqs2 overexpression does not affect bisexual syngamy (Figure S3A) but robustly stimulates different cellular differentiation events during the unisexual cycle. For instance, a strong enhancement of self-filamentation was observed in response to the increased Cqs2 expression (Figure S3B). Furthermore, the BMS analysis indicated a great increase in mature basidia in the P_*H3*_-*CQS2* strain compared with the XL280α strain, while the disruption of *CQS2* led to a significantly reduced population of mature basidia during unisexual development (Figure 3B; the P_*H3*_-*CQS2* strain: 95% CI: 0.14, 0.34, *P* = 5.33 × 10^−6^; the *cqs2*Δ mutant: 95% CI: −0.17, −0.03, *P* = 4.95 × 10^−3^, two-tailed Student’s *t*-test). In addition to these differentiation events, we showed the abundant expression of meiosis-specific Dmc1 in the P_*H3*_-*CQS2* strain at 3 days after unisexual induction when it was undetectable in most aerial hyphae in wild-type (Figure 3C). Lastly, an extremely robust unisexual sporulation was observed in response to Cqs2 overexpression, based on a method developed for the quantitative assessment of unisexual sporulation (see Materials and Methods for the details) (Figure 3D). Using this approach, we found that nearly 100% mini-colonies of the overexpression mutant produced hyphae with sporulated basidia at only 3 days after unisexual stimulation, when sporulation in the wild-type strain was extremely scarce (Figure 3D). A detailed phenotypic assay based on SEM detected ample basidia with four chains of basidiospores in the P_*H3*_-*CQS2* strain (Figure 3E), without evident difference in the morphology of basidiospores produced by XL280α and the overexpression strain (Figure 3E). Moreover, the germination of unisexual spores was not attenuated in the P_*H3*_-*CQS2* strain, which exhibited a germination rate even higher than that of the wild-type strain (Figure 3F), suggesting physiological normality in the spores derived from the strain with overexpression of Cqs2. Additionally, *CQS2* overexpression was able to largely restore the defect in the unisexual cycle to produce basidiospores in the *cqs1*Δ mutant, suggesting that Cqs2 can successfully drive the unisexual cycle even in the absence of Qsp1 (Figure S3C).

**Figure 3.**
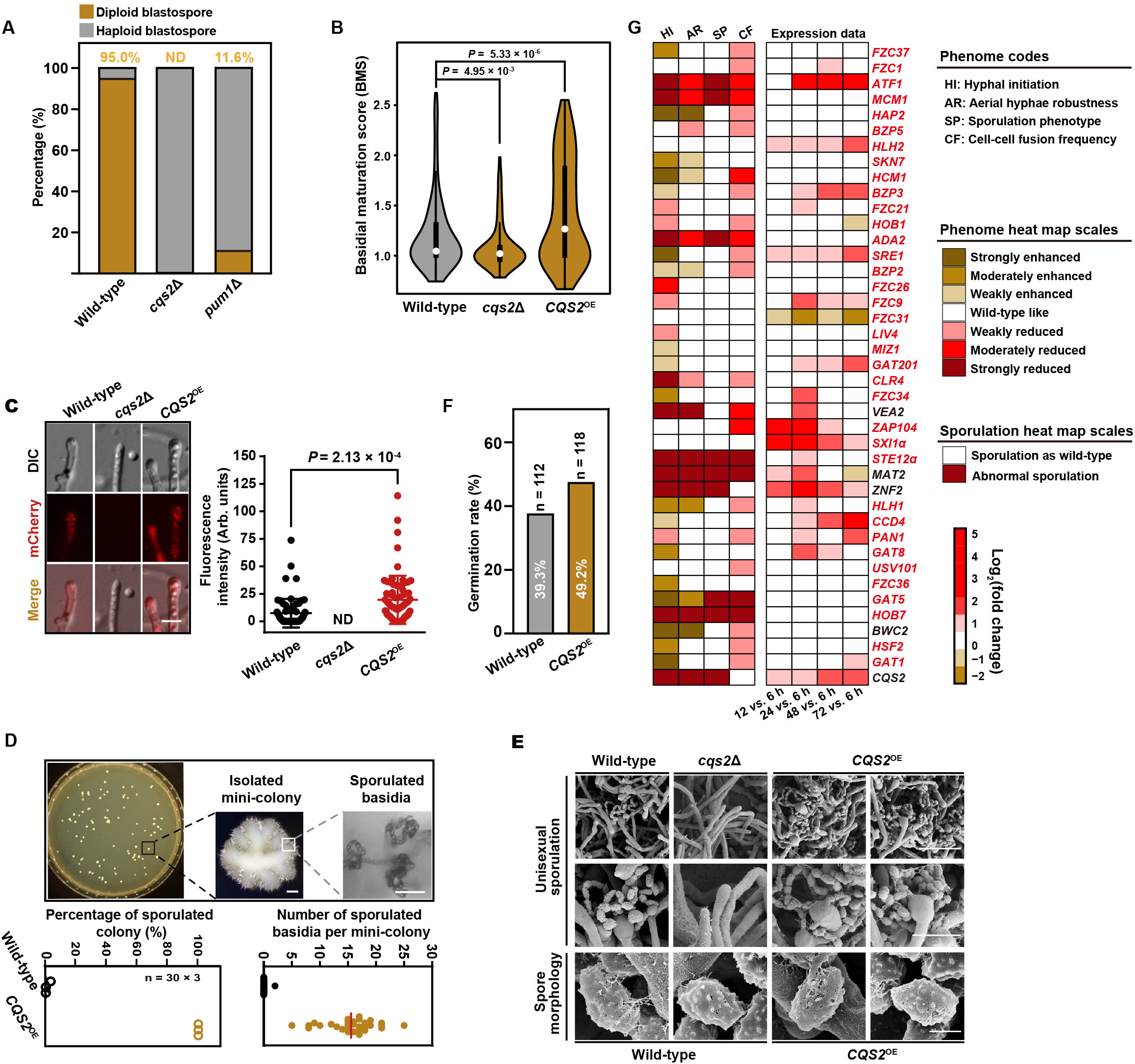
Cqs2 drives and coordinates multistage unisexual reproduction in C. *neoformans*. **(A)** FACS-based ploidy assessment of blastospores dissected from unisexual hyphae of different strains. For each strain, 40 blastospores dissected from different unisexual hyphae were tested for ploidy level. **(B)** Violin plot analysis showing BMS distribution of wild-type, *cqs2*Δ and *CQS2*^OE^ (P_*H3*_-*CQS2*). Hyphae with or without basidia were photographed at 5 days post unisexual stimulation and were randomly chosen for the BMS calculation (*N* = 150 for each strain). Two-tailed Student’s *t*-test. **(C)** The fluorescence-labeled strains were photographed at 3 days after inoculation on V8 agar to stimulate unisexual reproduction. For each strain, 50 basidia were examined for the expression of Dmc1-mCherry. Two-tailed Student’s *t*-test; scale bar, 10 μm. **(D)** Diagram showing the approach for quantitative analysis of unisexual sporulation (upper panel). Scale bars, 200 μm (upper middle panel), 20 μm (upper right panel). The overexpression of Cqs2 dramatically promoted the sporulation efficiency during unisexual reproduction. **(E)** The sporulation phenotypes of wild-type, *cqs2*Δ and *CQS2*^OE^. Scale bar, 10 μm (upper panel), 5 μm (middle panel), 1 μm (bottom panel). **(F)** Germination rate of unisexual basidiospores from *CQS2*^OE^ compared to wild-type. Basidiospores were dissected from strains cultured on V8 agar for 7 days. **(G)** Phenotype scores are represented in distinct colors based on qualitative or semi-quantitative phenotypic analysis of >2 independent TF mutant strains (left panel). Phenotype strengths (weak, intermediate and strong) are distinguished in gradients of red or yellow, as indicated in the right panel. Measurement of hyphal initiation (filamentation frequency) and cell-cell fusion efficiency based on unilateral mating was performed as described in Materials and Methods. Genes reported previously to control unisexual development are denoted in black, whereas genes that we newly deleted in XL280α are denoted in red. Transcriptional dynamics for each TF during unisexual development were based on the RNA sequencing data reported in the previous XL280α sexual reproduction paper [16].

We next ask whether there exist other TFs that likewise exert specific function on the promotion and coordination of the multistage unisexual cycle but are not important for bisexual syngamy. To address this, we performed the phenotypic assays in 41 TF deletion strains in the XL280α background. These TFs have been previously documented to regulate sexual development in *Cryptococcus neoformans* species complex. They include 5 regulators reported to activate unisexual filamentation in the serotype D XL280α background [33, 39] and 36 TFs previously identified to positively or negatively affect morphogenesis during bisexual reproduction in a systematic study targeting the serotype A H99 strain [41], which is unable to undergo solo unisexual reproduction. Consequently, the coding genes of these 36 TFs were deleted in XL280α in order to examine their function on unisexual development. Based on the phenotypic evaluation of all 41 mutants, 38 TFs (~92.7%) were shown to affect at least one stage of unisexual development (hyphal initiation, aerial morphogenesis and sporulation) or bisexual syngamy (Figures 3G). Among the mutants, only *clr4*Δ and *znf2*Δ displayed reduced filamentation with mildly decreased or normal cell-cell fusion ability that resemble the phenotypes observed in the *cqs2*Δ mutant (Figures 3G). However, unlike *CQS2*, overexpression of either *ZNF2* or *CLR4* were shown to be not sufficient to promote and accomplish the multistage unisexual reproduction but severely impaired the late developmental stages, including basidial formation and sporulation, as illustrated in Figure S4.

### Pum1, as a key target of Cqs2, governs unisexual autopolyploidization

To explore the molecular basis underlying the pivotal function mediated by Cqs2 in the unisexual cycle, we employed two independent Chromatin Immunoprecipitation Sequencing (ChIP-seq) experiments (Figure 4A). The experiments were performed using anti-mCherry antibody and anti-FLAG antibody against the strains, which harbor the genes encoding mCherry-tagged and FLAG-tagged Cqs2, respectively. Two ChIP-seq approaches co-identified significant overlapping of binding events (Figures 4 A and 4B; Table S3; *P* = 5.32 × 10^−42^), which were further analyzed to determine the accurate direct regulon of Cqs2 and its binding motif during unisexual reproduction (Figure 4C). Based on the ChIP-seq analysis, Cqs2 does not directly bind to the promoter regions of any sex-determination genes (Figure 4D), again supporting its dispensable function on syngamy that strictly requires sex-determination system. Expectedly, the direct regulon of Cqs2 includes multiple genes involved in various stages of the unisexual cycle, including filamentation, basidial formation, meiosis and sporulation (Figure 4D). Among these genes, we focused on *PUM1*, which encodes a Pumilio family RNA-binding protein, due to that it, like Cqs2, was reported to be similarly involved in the coordination of multiple sexual differentiation events but is not essential for bisexual syngamy [36]. We further confirmed this idea on the basis of the following findings: i) ChIP-seq together with qRT-PCR assays indicated that Cqs2 stimulates the expression of *PUM1* during unisex in a direct manner (Figures 4E and 4F); ii) transcriptomically, ~45.4% targets of Cqs2 were shared in the Pum1 regulon (Figure 4G and Table S4, *P* = 3.42 × 10^−6^, Fisher’s exact test); iii) the disruption of *PUM1* led to a severe defect of unisex-specific endoreplication and dramatically reduced the production of diploid blastospores during unisex compared with wild-type (Figure 3A); and iv) Cqs2 overexpression in the *pum1*Δ mutant failed to restore the defects of unisexual differentiation (Figures 5A and 5B). Furthermore, we found that the overexpression of Pum1 is sufficient to drive autopolyploidization, even under sex-suppressing conditions (Figures 5C and 5D). Once Pum1 was forced to express, a considerable increase in cell size was detected in a large population of cells with a high ploidy state (Figures 5C and 5D). Notably, the cells with high ploidy retained yeast morphology (Figures S5A-S5C), suggesting that Pum1-activated autopolyploidization can be uncoupled with self-filamentation. These evidences support that Pum1 is a vital factor for inducing autopolyploidization, and its induction occurs regardless of the conditions and developmental cues. We further showed that Pum1 overexpression can efficiently stimulate autopolyploidization in the absence of Cqs2 but not *vice versa* (Figure S5D). These results suggest that Pum1 activates self-diploidization during unisex as a key target of Cqs2 (Figure S5E).

**Figure 4.**
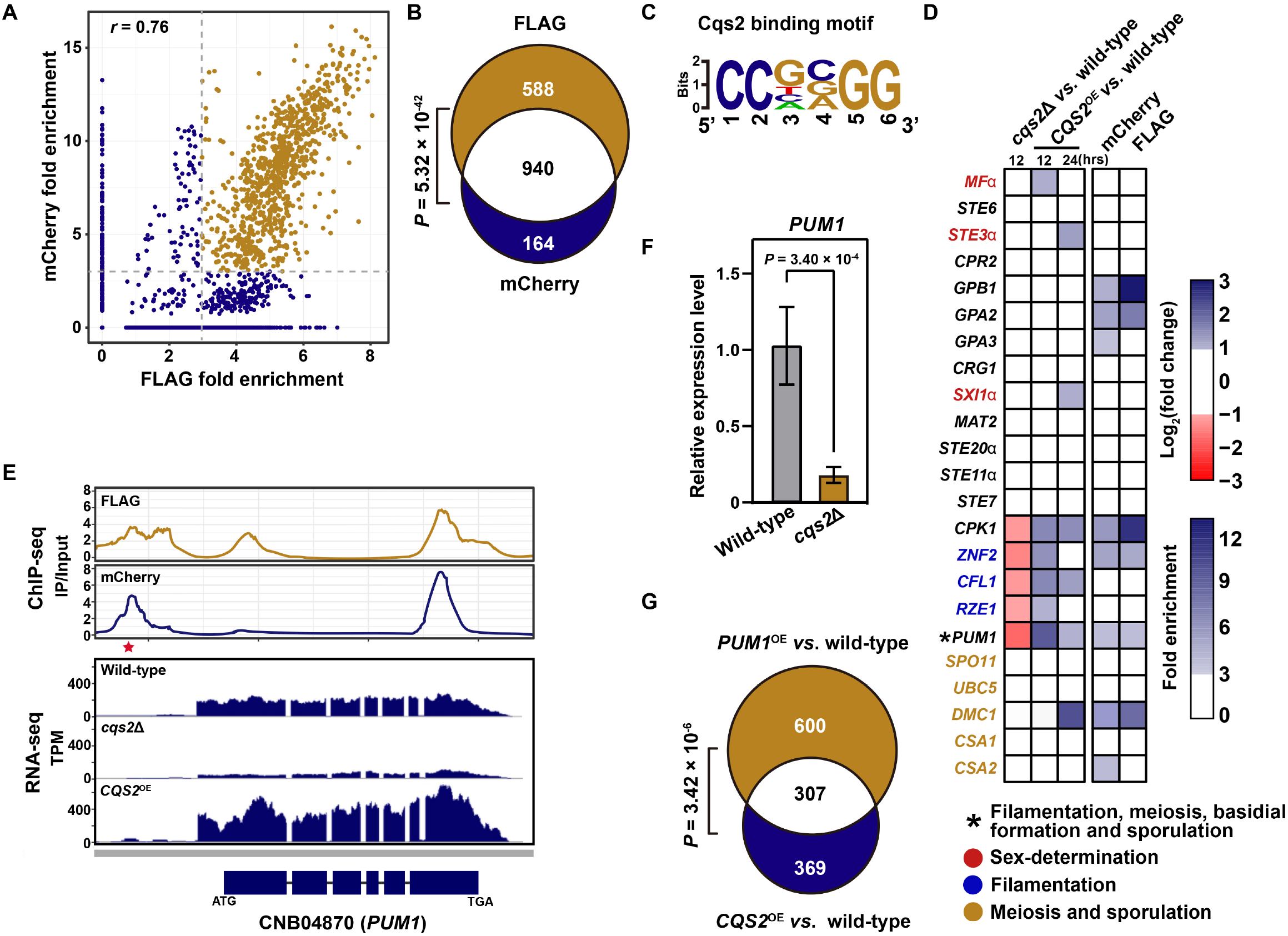
Cqs2 governs unisex by coordinating the expression of multiple genes involved in various unisexual stages. **(A)** Binding events identified by FLAG and mCherry tagged ChIP-seq approaches, respectively. **(B)** Significant overlap between binding events identified by two ChIP-seq approaches. **(C)** The binding motif of Cqs2. **(D)** Cqs2 directly activates the genes involved in various stages during sexual development but cannot bind to the promoters of the sex-determination genes. **(E)** Cqs2 controls the expression of *PUM1* in a direct manner. The asterisk indicates the Cqs2 binding site at the promoter of *PUM1*. **(F)** qRT-PCR showing the mRNA level of *PUM1* in *cqs2*Δ during unisexual reproduction compared to wild-type. Data are presented as the mean ± SD of six independent experiments, two-tailed Student’s *t*-test. **(G)** The overlap between *CQS2*-regulated genes revealed by RNA-seq analysis of the *CQS2* overexpression strain and *PUM1*-controlled genes, which were identified based on the published RNA-seq data targeting the *PUM1*^OE^ strain [16]. Fisher’s exact test.

**Figure 5.**
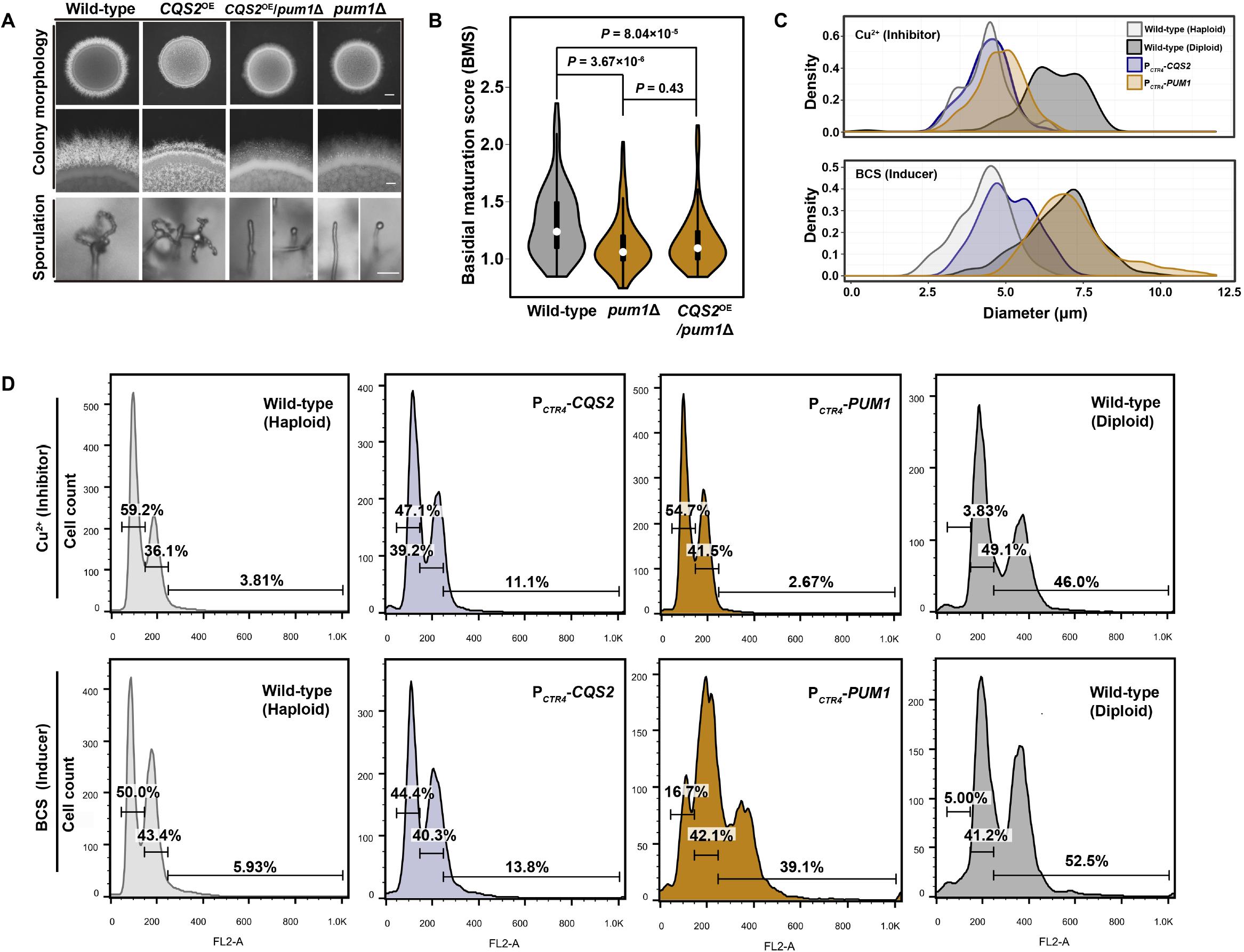
Pum1 is a key target of Cqs2 during unisexual autopolyploidization. **(A)** The colony morphology, self-filamentation and sporulation phenotypes of various mutants. Hyphae and chains of basidiospores were photographed after 7 days of growth on V8 medium. Scale bars, 1 mm (upper and middle panel), 20 μm (bottom panel). **(B)** Overexpression of *CQS2* cannot restore the defect of basidial maturation in the *pum1*Δ mutant. Strains were cultured on V8 agar for 7 days, and 150 hyphae of each strain were randomly chosen for BMS calculation. Two-tailed Student’s *t*-test. **(C)** Cell size distribution of wild-type, P_*CTR4*_-*PUM1* and P_*CTR4*_-*CQS2* cultured under sex-suppressing condition (YPD liquid medium). Strains were cultured in YPD liquid medium for 3 days in the presence of copper (inhibitor of *CTR4* promoter, final concentration 25 μM) or BCS (inducer of *CTR4* promoter, final concentration 20 μM) and over 150 cells of each strain were examined for their diameters. **(D)** FACS-based ploidy level assessment of wild-type, P_*CTR4*_-*PUM1* and P_*CTR4*_-*CQS2* cultured under sex-suppressing condition. Strains were cultured in YPD liquid medium for 3 days in the presence of copper (final concentration 25 μM) or BCS (final concentration 20 μM), and the ploidy of each strain was subsequently assessed by FACS analysis.

## Discussion

Meiotic reproduction is not the main way for proliferation in most fungal species, including *C. neoformans*, but acts as an adaptation strategy in response to external stress [42]. Consistently, fungal sex is normally induced under stress conditions [42]. The benefit generated through sexual reproduction in fungi is considered to offer the genetic diversity among sexual progenies, which is created by meiotic recombination between parental genomes *via* syngamy and maximizes fungal lineage and community fitness in fluctuating environments [6, 43–46]. However, the syngamy-independence of unisex in *C. neoformans* results in an extreme inbreeding feature. This may suggest that *C. neoformans* needs unisexual reproduction for unique advantages other than syngamy-dependent genetic variance in sexual offspring. Indeed, a considerable dispersal benefit has been proven to be related to unisexual air-borne spores and filaments [47], which also help prevent against engulfment by amoeboid predator [22]. In addition, unisexual reproduction can produce ploidy variance among progenies that contributes to *de novo* genotypic and phenotypic plasticity [48].

A homeobox family regulator, pheromone and pheromone receptor represent the conserved sex-determination components across species belonging to Basidiomycota. In basidiomycetes, the recognition between pheromone and its cognate receptor provides the key compatibility checkpoint that underlies syngamy during heterothallic and homothallic mating [49]. Once mating partners with compatible pheromone and cognate receptor pairs have fused, homeobox family regulators regulate a transcriptional cascade that promotes further sexual development [40, 49, 50]. *C. neoformans* α strains contain an unusual *MAT* locus that is over 100 kb in size and contains approximately two dozen genes [51]. In this locus, five genes are sufficient to specify the α cell type, including the three paralogue genes encoding Mfα as well as the two genes, which produce the GPCR pheromone receptor Ste3α and the homeobox regulator Six1α, respectively [29]. Among these α identity factors, the pheromone Mfα and the pheromone receptor Ste3α appear to play the central role in the establishment of sexual identity. Replacement of the coding genes of pheromones and the pheromone receptor in the α strain with the analogous genes from the **a** strain leads to the resulting strain behaving like the **a** strain [29]. It has been demonstrated that the perception of the fungal pheromone by its compatible receptor is the key event before syngamy being decided [52]. Syngamy-independent characteristics of α unisexual development may be because of the incompatibility of Mfα and Ste3α during unisex. This potential incompatibility likely provides a molecular basis to avoid the co-occurrence of syngamy and autopolyploidization that potentially causes an inappropriate ploidy state during unisex in *C. neoformans*.

It has been proposed by independent laboratories that meiosis was an adaptive response to auto-polyploidy potentially caused by endoreplication or endomitosis in early eukaryotes, and syngamy might evolve later [53, 54]. In this regard, meiosis evolved as a mechanism to correct for spontaneous whole-genome duplication or to enable ploidy cycling to maximize ploidy state-related fitness, independent of any role in shuffling genetic composition by recombination [53]. Considering the essential role of sex-determination program in syngamy (fertilization) in modern eukaryotic species [3, 55], the early eukaryotes that undergo syngamy-dispensable sexual reproduction may lack sex-determination system. Our results provide the direct evidence of sexual reproduction independent of the sex-determination system in the modern fungus *C. neoformans*. We determined that a non-cell-identity factor Cqs2 as the critical regulatory determinant activates unisex-specific self-diploidization and orchestrates various stages and therefore safeguards the formation of unisexual meiospore progenies. Conversely, Cqs2 plays a marginal role in the control of bisexual syngamy, which is dominated by the sex-determination system. ChIP-seq analysis confirmed that Cqs2 binds to the upstream regions of the genes involved in various unisexual phases but not those of the sex-determination genes. Phylogenetic analysis indicated that Cqs2 orthologs from different species belonging to *Cryptococcaceae* share high similarity in the protein sequence (greater than 40% identity and over 95% coverage) (data not shown). This similarity suggests a possibility that the syngamy-independent unisexual life cycle may be widespread in this fungal clade, given the pivotal role of Cqs2 in this reproduction mode.

## Materials and Methods

### Strains, media, and growth conditions

The strains used in this study are listed in Table S5. Yeast cells were grown routinely at 30°C on YPD (1% yeast extract, 2% Bacto peptone, 2% dextrose, and 2% Bacto agar) medium. Unisexual differentiation and bisexual cell-cell fusion assays were conducted on V8 agar (0.5 g/liter KH_2_PO_4_, 4% Bacto agar and 5% V8 juice, pH = 7.0) at 25°C in the dark for the designated time period. For dissection of blastospores, strains were cultured on Murashige and Skoog (MS) medium minus sucrose (Sigma-Aldrich) for 2 weeks at 25°C in the dark.

### Filamentation assay, sporulation assay and BMS assay

Quantitative analysis of filamentation frequency was conducted as previously described [39]. Briefly, low-density cells were dropped onto V8 agar to allow the formation of isolated mini-colonies after 24 hours of cultivation. The mini-colonies exhibited heterogeneity in filamentation, and the strength of unisexual induction is reflected by the filamentous incidence among mini-colonies. Unisexual reproduction was examined microscopically for the formation of sexual hyphae and chains of basidiospores. For quantitative analysis of sporulation, the cells of different strains were plated onto V8 medium at a low cell density to form isolated mini-colonies. After one-week of culturing, the aerial hyphae residing in mini-colonies showed heterogeneity in differentiation into sporulated basidia, and the sporulation incidence among the hyphae from the mini-colonies positively correlates with the strength of unisexual sporulation. The BMS analysis was performed as previously described [16].

### Microscopy and fluorescence

To examine the expression and subcellular localization of Dmc1, strains harboring P_*DMC1*_-*DMC1*-*mCherry* were spotted onto V8 agar at 25°C in the dark for designated period of time. Images were taken and performed with a Zeiss Axioplan 2 imaging system with the AxioCam MRm camera software Zen 2011 (Carl Zeiss Microscopy).

### Basidiospore germination assay

Cells (original absorbance *A*_600_ = 1.0) were spotted onto V8 agar for 7 days at 25°C in the dark, and basidiospores were dissected on YPD medium using a SporePlay dissection microscope (Singer Instruments). The dissected spores were incubated at 30°C for 3-5 days until colonies were formed. The number of visible colonies formed by germinated spores was counted. The basidiospore germination rate was determined as a ratio of the number of visible colonies to the total number of basidiospores dissected.

### Cell-cell fusion assay

Cell-cell fusion assay was performed as previously described [28]. The strains were grown on YPD plates for 2 days. The cells were collected and washed twice with ddH_2_O and diluted to a final *A*_600_ = 1.0. Equal numbers of mating partners were mixed, and 10 microliters of the mixture were spotted onto V8 agar. After incubation for 15 hours in the dark at 25°C, the cells were then removed, washed with ddH_2_O, and fused products were selected on YPD medium supplemented with hygromycin and G418. CFUs (colony-forming units) were counted to measure the efficiency of cell fusion.

### Blastospore dissection and flow cytometry

Blastospore dissection was preformed according to a previous study [25]. Cells (original absorbance *A*_600_ = 1.0) were dropped onto MS agar for 2 weeks at 25°C in the dark, and blastospores were dissected on YPD medium using a SporePlay dissection microscope (Singer Instruments). The ploidy of the blastospores was determined by fluorescence activated cell sorting (FACS) analysis as previously described [56].

Briefly, cells were harvested after growth on a YPD plate, washed in phosphate buffered saline (PBS) and fixed in 70% ethanol at 4°C overnight. The fixed cells were washed once with 1 ml NS buffer (10 mM Tris-HCl pH 7.6, 250 mM sucrose, 1 mM EDTA, 1 mM MgCl_2_, 0.1 mM CaCl_2_, 0.1 mM ZnCl_2_, 0.4 mM phenylmethyl sulphonyl fluoride, 7 mM mercaptoethanol), and then stained with propidium iodide (10 mg/ml) in 200 μl of NS buffer containing RNaseA (1 mg/ml) at 4°C overnight. Stained cells were then diluted into Tris-HCl (pH = 8.0) buffer and sonicated. A total of 10,000 cells were analyzed for each sample, using the FL2 channel on a BD FACSCalibur™ at the Beijing Regional Center of Life Science Instrument, Chinese Academy of Science. FlowJo software was used to perform data analysis.

### Gene knockout and overexpression

Targeted gene deletion was performed as described previously [57]. Briefly, the deletion construct of ~1 kb flanking the target gene and splitting part of NEO (Neomycin)/ NAT (Nourseothricin) resistance cassette was introduced into relevant *Cryptococcus* strains by biolistic transformation. The α-null strain was constructed by *TRACE* [58]. All mutants generated were genetically and transcriptionally confirmed by PCR and qRT-PCR, respectively. For *CQS2* overexpression, the coding-region of *CQS2* was placed downstream of the promoter of histone H3 (using promoter swap), whose expression was abundant and steady throughout unisexual development, based on reanalyzing previous RNA-seq data [16]. For the inducible expression of *PUM1* and *CQS2*, the overexpression constructs were created by amplifying the entire ORFs and inserting into plasmid pXC-mCherry after the *CTR4* promoter as previously described [57]. The overexpression constructs were linearized and introduced into relevant *Cryptococcus* strains by electroporation. The primers used for the generation of the mutants are listed in Table S6.

### Scanning electron microscopy (SEM)

SEM was performed at the Beijing Regional Center of Life Science Instrument, Chinese Academy of Science. Samples were prepared as previously described [25]. The cells were spotted onto V8 solid medium at 25°C in the dark for 7 days. Unisexual colony patches were then excised and fixed in phosphate-buffered glutaraldehyde (pH 7.2) at 4°C for more than 2 hours. The fixed samples were rinsed in deionized distilled water (ddH_2_O) three times for 6 min, 7 min and 8 min. The samples were dehydrated in a graded ethanol series (50%, 70%, 85% and 95%) for 14 min for each concentration and dehydrated in 100% ethanol three times for 15 min. After dehydration, the blocks were critical-point dried with liquid CO_2_ (Leica EM CPD300, Germany) and sputter coated with gold-palladium (E-1045 ion sputter, Hitachi, Japan). The samples were viewed and images were taken using a Quanta200 scanning electron microscope (FEI, USA).

### RNA purification and quantitative RT-PCR analysis

RNA extraction and quantitative real-time PCR (qRT-PCR) were carried out as described previously [39]. Briefly, the cells were frozen in liquid nitrogen and ground to a fine powder, total RNA was extracted and purified using an Ultrapure RNA Kit (Kangweishiji, CW0581S) according to the manufacturer’s instructions. A Fastquant RT Kit (Tiangen KR106-02, with gDNase) was used for first-strand cDNA synthesis following the manufacturer’s instructions. qRT-PCR was conducted using power SYBR qPCR premix reagents (KAPA) in a CFX96 Touch™ Real-time PCR detection system (BioRad), and relative transcript levels were calculated to fold change and normalized to the housekeeping gene *TEF1* using the comparative *C*_*t*_ method as previously described [57]. The primers used for qRT-PCR are listed in Table S6.

### RNA-seq and data analysis

For RNA-seq analysis in this study, the different strains were cultured in YPD liquid medium at 30°C overnight. The cells were washed with ddH_2_O and spotted on V8 medium to stimulate unisexual reproduction. The RNA level and integrity of each sample were evaluated using a Qubit RNA Assay Kit on a Qubit 2.0 Flurometer (Life Technologies, CA, USA) and RNA Nano 6000 Assay Kit with the Bioanalyzer 2100 system (Agilent Technologies, CA, USA), respectively. RNA purity was assessed using a Nano Photometer spectrophotometer (IMPLEN, CA, USA). The transcriptome libraries were generated using VAHTS mRNA-seq v2 Library Prep Kit (Vazyme Biotech Co., Ltd, Nanjing, China) according to the manufacturer’s instructions. Sequencing the transcriptome libraries was performed by Annoroad Gene Technology Co., Ltd (Beijing, China). The samples were clustered using VAHRS RNA adapters set1/set2 and sequenced on an Illumina platform. For RNA-seq analysis, the quality of sequenced clean data was performed using FastQC v0.11.5 software. As a result, ~2 GB of clean data for each sample were mapped to the genome sequence of *C. neoformans* XL280α using STAT_2.6.0c. The gene expression level was measured in TPM by Stringtie v1.3.3 to determine unigenes. All unigenes were subsequently aligned against the well-annotated genome of JEC21 (which is congenic to JEC20**a** that served as the parent strain to generate XL280α through a cross with B3501α). The differential expression of genes (DEGs) was assessed using DEseq2 v1.16.1 of the R package and defined based on the fold change criterion (log2|fold-change| > 1.0, q value < 0.01). In all RNA-seq assays performed in this study, three biological replicates were included.

### Chromatin Immunoprecipitation Sequencing (ChIP-seq) and data analysis

ChIP was performed as described previously with modifications [59]. Briefly, strains expressing Cqs2-Flag and Cqs2-mCherry were diluted and spotted onto V8 agar containing 200 μM BCS for 8 hours at 25°C in the dark. The cells were harvested and cross-linked with 1% formaldehyde at room temperature for 15 min. Glycine (2.5 M) was then added to a final concentration of 125 mM, and the treated cultures were incubated for another 5 min at room temperature. Cell pellets were harvested by centrifugation at 4000 rpm for 5 min at 4°C and washed three times with ice-cold water. Cell pellets were then ground to a fine powder in liquid nitrogen. Chromatin fragmentation was performed by digestion using 2 units of microccocal nuclease for 15 min at 37°C. The nuclear membrane was disrupted by sonication in a Bioruptor water bath sonicator (Diagenode, 30 s on, 30 s off, high setting, 3 cycles) at 4°C. After centrifugation at 12,000g for 10 min at 4 °C, the supernatant was collected, and a 50 μl aliquot of sample was reserved as ChIP input material. For the strain samples harboring Cqs2 tagged with mCherry or FLAG, 20 μl of RFP-TRAP magnetic beads (Chromo Tek) or anti-FLAG M2 magnetic beads (SIGMA-Aldrich) were added to the remaining lysate. All samples were incubated overnight at 4°C with agitation. The beads were washed 5 min at 4 °C with ice-cold buffers as follows: twice with lysis buffer, once with high–salt buffer, once with LiCl buffer, and once with TE buffer. After the final wash, the bound chromatin was eluted with 250 μl elution buffer. Cross-links of input and IP material were reversed by adding 10 μl 5 M NaCl and 4 μl 10 mg ml^−1^ Proteinase K (Promega), and incubated for 4 hours at 65°C. DNA was extracted using phenol-chloroform and ethanol precipitation. The isolated DNA was dried and resuspended in 20 μl TE containing 10 μg ml^−1^ RNase. For library construction, 5 ng of ChIP input and IP material were end-repaired with Klenow DNA polymerase, A-tailed using Klenow (exo-) and ligated to adapters. The ligated fragments were amplified using indexed primers (Illumina) for 8-10 PCR cycles. DNA was purified with 1.8 volume Ampure XP DNA purification beads between each step. Library sequencing was performed on an Illumina Hi-Seq 2000. The raw data were aligned using Bowtie2 (version 2.1.0) [60] with default parameters. Indexed and sorted bam files of each dataset were created using SAMtools (version 1.5) [61]. ChIP-Seq peaks were called using MACS2 (version 2.1.1.20160309) [62]. Annotation used ChIPseeker (R 1,12,1) to locate each peak to promoter regions of related genes. Chromosomal traces were plotted using Integrative Genomics Browser 9.0.0 (Broad Institute). ChIP-Seq peaks of the mCherry and FLAG tags were merged using HOMER (4.9.1), and 50 bp upstream and downstream merged peaks were selected to find the conserved binding motif (HOMER 4.9.1). The motifs were generated for widths of 6-14 bps for both strands represented by the sequences, and the length of the motif reported was determined by the lowest *P*-value.

### Statistical analyses

Statistical analyses were performed using R, version 3.4.2. We used a two-tailed unpaired Student’s *t*-test to compare the mean florescence intensity, basidial maturation score or transcript levels between two groups. Fisher’s exact test and chi-square test were utilized to evaluate the significance between two sets of genes. Permutation test was utilized for analysis of the significance of GSEA enrichment. A two-sided *P* < 0.05 was considered significant, and *P* <0.001 were considered very significant. The data are shown as means ± SD from three or more independent experiments.

## Acknowledgements

We would like to thank Dr. Joseph Heitman, Dr. Alexander Idnurm, Dr. Chaoyang Xue, Dr. Xiaorong Lin, Dr. Sheng Sun, Dr. Ci Fu and Dr. Youbao Zhao for critical reading and helpful suggestions. We thank Dr. Ci Fu for important advice about *C. neoformans* blastospore dissection.

## Funding

This work was financially supported by the Key Research Program of the Chinese Academy of Sciences (QYZDB-SSW-SSMC040), the National Science and Technology Major Project (2018ZX10101004), the National Natural Science Foundation of China (Grants 31622004, 31570138 and 31770163). Linqi Wang is a member of the “Thousand Talents Program”.

## Author contributions

All authors contributed to the data analysis. P.H., H.L., L.C., G.-J.H., X. T., X.Y., C.Z., C.L., C.T., E.Y. and L.W. designed the experiments. P.H. and H.L. conducted most of the studies. L.C., C.T., and E.Y. conducted most of the bioinformatics assays. X. T. conducted most of the RNA extraction and qRT-PCR. G.-J.H., H.L, and P.H. construct most of the strains and conducted the phenotypic assays. H.L. and C.L. performed the scanning electron microscopy experiments. P. H., H.L, and C.Z. conducted most of FACS analysis. X.Y. conducted the basidiospore dissection. E.Y., and L.W. contributed reagents/materials/analysis tools. P.H., H.L., L. C., X. T. and L.W. wrote the paper with contributions from all other authors.

## Competing interests

The authors declare that they have no competing interests.

## Data and materials availability

All data needed to evaluate the conclusions in the paper are present in the paper and/or the Supplementary Materials. The sequencing data has been deposited at the Gene Expression Omnibus (GEO) with accession number GSE135566. All data supporting the findings of this study are available from the corresponding author upon request.

## References

1. Fraser, J.A., and Heitman, J. (2005). Chromosomal sex-determining regions in animals, plants and fungi. Curr Opin Genet Dev 15, 645–651.

2. David-Palma, M., Sampaio, J.P., and Goncalves, P. (2016). Genetic Dissection of Sexual Reproduction in a Primary Homothallic Basidiomycete. PLoS Genet 12, e1006110.

3. Jones, S.K., Jr., and Bennett, R.J. (2011). Fungal mating pheromones: choreographing the dating game. Fungal Genet Biol 48, 668–676.

4. Neaves, W.B., and Baumann, P. (2011). Unisexual reproduction among vertebrates. Trends Genet 27, 81–88.

5. Rajasingham, R., Smith, R.M., Park, B.J., Jarvis, J.N., Govender, N.P., Chiller, T.M., Denning, D.W., Loyse, A., and Boulware, D.R. (2017). Global burden of disease of HIV-associated cryptococcal meningitis: an updated analysis. Lancet Infect Dis 17, 873–881.

6. Idnurm, A., Bahn, Y.S., Nielsen, K., Lin, X., Fraser, J.A., and Heitman, J. (2005). Deciphering the model pathogenic fungus Cryptococcus neoformans. Nat Rev Microbiol 3, 753–764.

7. Heitman, J., Carter, D.A., Dyer, P.S., and Soll, D.R. (2014). Sexual reproduction of human fungal pathogens. Cold Spring Harb Perspect Med 4.

8. Fraser, J.A., Giles, S.S., Wenink, E.C., Geunes-Boyer, S.G., Wright, J.R., Diezmann, S., Allen, A., Stajich, J.E., Dietrich, F.S., Perfect, J.R., et al. (2005). Same-sex mating and the origin of the Vancouver Island Cryptococcus gattii outbreak. Nature 437, 1360–1364.

9. Voelz, K., Ma, H., Phadke, S., Byrnes, E.J., Zhu, P., Mueller, O., Farrer, R.A., Henk, D.A., Lewit, Y., Hsueh, Y.P., et al. (2013). Transmission of Hypervirulence traits via sexual reproduction within and between lineages of the human fungal pathogen cryptococcus gattii. PLoS Genet 9, e1003771.

10. Sun, S., Billmyre, R.B., Mieczkowski, P.A., and Heitman, J. (2014). Unisexual reproduction drives meiotic recombination and phenotypic and karyotypic plasticity in Cryptococcus neoformans. PLoS Genet 10, e1004849.

11. Giles, S.S., Dagenais, T.R., Botts, M.R., Keller, N.P., and Hull, C.M. (2009). Elucidating the pathogenesis of spores from the human fungal pathogen Cryptococcus neoformans. Infect Immun 77, 3491–3500.

12. Velagapudi, R., Hsueh, Y.P., Geunes-Boyer, S., Wright, J.R., and Heitman, J. (2009). Spores as infectious propagules of Cryptococcus neoformans. Infect Immun 77, 4345–4355.

13. Botts, M.R., and Hull, C.M. (2010). Dueling in the lung: how Cryptococcus spores race the host for survival. Curr Opin Microbiol 13, 437–442.

14. Kozubowski, L., and Heitman, J. (2012). Profiling a killer, the development of Cryptococcus neoformans. FEMS Microbiol Rev 36, 78–94.

15. Ballou, E.R., and Johnston, S.A. (2017). The cause and effect of Cryptococcus interactions with the host. Curr Opin Microbiol 40, 88–94.

16. Liu, L., He, G.J., Chen, L., Zheng, J., Chen, Y., Shen, L., Tian, X., Li, E., Yang, E., Liao, G., et al. (2018). Genetic basis for coordination of meiosis and sexual structure maturation in Cryptococcus neoformans. Elife 7.

17. Huang, M., and Hull, C.M. (2017). Sporulation: how to survive on planet Earth (and beyond). Curr Genet 63, 831–838.

18. Kwon-Chung, K.J. (1976). Morphogenesis of Filobasidiella neoformans, the sexual state of Cryptococcus neoformans. Mycologia 68, 821–833.

19. Lin, X., Hull, C.M., and Heitman, J. (2005). Sexual reproduction between partners of the same mating type in Cryptococcus neoformans. Nature 434, 1017–1021.

20. Ni, M., Feretzaki, M., Sun, S., Wang, X., and Heitman, J. (2011). Sex in fungi. Annu Rev Genet 45, 405–430.

21. Fu, C., Sun, S., Billmyre, R.B., Roach, K.C., and Heitman, J. (2015). Unisexual versus bisexual mating in Cryptococcus neoformans: Consequences and biological impacts. Fungal Genet Biol 78, 65–75.

22. Zhao, Y., Lin, J., Fan, Y., and Lin, X. (2019). Life Cycle of Cryptococcus neoformans. Annu Rev Microbiol.

23. Wang, L., and Lin, X. (2011). Mechanisms of unisexual mating in Cryptococcus neoformans. Fungal Genet Biol 48, 651–660.

24. Roach, K.C., Feretzaki, M., Sun, S., and Heitman, J. (2014). Unisexual reproduction. Adv Genet 85, 255–305.

25. Fu, C., and Heitman, J. (2017). PRM1 and KAR5 function in cell-cell fusion and karyogamy to drive distinct bisexual and unisexual cycles in the Cryptococcus pathogenic species complex. PLoS Genet 13, e1007113.

26. Wang, L., Tian, X., Gyawali, R., and Lin, X. (2013). Fungal adhesion protein guides community behaviors and autoinduction in a paracrine manner. Proc Natl Acad Sci U S A 110, 11571–11576.

27. McClelland, C.M., Chang, Y.C., Varma, A., and Kwon-Chung, K.J. (2004). Uniqueness of the mating system in Cryptococcus neoformans. Trends Microbiol 12, 208–212.

28. Lin, X., Jackson, J.C., Feretzaki, M., Xue, C., and Heitman, J. (2010). Transcription factors Mat2 and Znf2 operate cellular circuits orchestrating opposite- and same-sex mating in Cryptococcus neoformans. PLoS Genet 6, e1000953.

29. Stanton, B.C., Giles, S.S., Staudt, M.W., Kruzel, E.K., and Hull, C.M. (2010). Allelic exchange of pheromones and their receptors reprograms sexual identity in Cryptococcus neoformans. PLoS Genet 6, e1000860.

30. Chung, S., Karos, M., Chang, Y.C., Lukszo, J., Wickes, B.L., and Kwon-Chung, K.J. (2002). Molecular analysis of CPRalpha, a MATalpha-specific pheromone receptor gene of Cryptococcus neoformans. Eukaryot Cell 1, 432–439.

31. Shen, W.C., Davidson, R.C., Cox, G.M., and Heitman, J. (2002). Pheromones stimulate mating and differentiation via paracrine and autocrine signaling in Cryptococcus neoformans. Eukaryot Cell 1, 366–377.

32. Gyawali, R., Zhao, Y., Lin, J., Fan, Y., Xu, X., Upadhyay, S., and Lin, X. (2017). Pheromone independent unisexual development in Cryptococcus neoformans. PLoS Genet 13, e1006772.

33. Feretzaki, M., and Heitman, J. (2013). Genetic circuits that govern bisexual and unisexual reproduction in Cryptococcus neoformans. PLoS Genet 9, e1003688.

34. Zhai, B., Zhu, P., Foyle, D., Upadhyay, S., Idnurm, A., and Lin, X. (2013). Congenic strains of the filamentous form of Cryptococcus neoformans for studies of fungal morphogenesis and virulence. Infect Immun 81, 2626–2637.

35. Gyawali, R., Upadhyay, S., Way, J., and Lin, X. (2017). A Family of Secretory Proteins Is Associated with Different Morphotypes in Cryptococcus neoformans. Appl Environ Microbiol 83.

36. Wang, L., Tian, X., Gyawali, R., Upadhyay, S., Foyle, D., Wang, G., Cai, J.J., and Lin, X. (2014). Morphotype transition and sexual reproduction are genetically associated in a ubiquitous environmental pathogen. PLoS Pathog 10, e1004185.

37. Lee, H., Chang, Y.C., Nardone, G., and Kwon-Chung, K.J. (2007). TUP1 disruption in Cryptococcus neoformans uncovers a peptide-mediated density-dependent growth phenomenon that mimics quorum sensing. Mol Microbiol 64, 591–601.

38. Homer, C.M., Summers, D.K., Goranov, A.I., Clarke, S.C., Wiesner, D.L., Diedrich, J.K., Moresco, J.J., Toffaletti, D., Upadhya, R., Caradonna, I., et al. (2016). Intracellular Action of a Secreted Peptide Required for Fungal Virulence. Cell Host Microbe 19, 849–864.

39. Tian, X., He, G.J., Hu, P., Chen, L., Tao, C., Cui, Y.L., Shen, L., Ke, W., Xu, H., Zhao, Y., et al. (2018). Cryptococcus neoformans sexual reproduction is controlled by a quorum sensing peptide. Nat Microbiol 3, 698–707.

40. Hull, C.M., Boily, M.J., and Heitman, J. (2005). Sex-specific homeodomain proteins Sxi1alpha and Sxi2a coordinately regulate sexual development in Cryptococcus neoformans. Eukaryot Cell 4, 526–535.

41. Jung, K.W., Yang, D.H., Maeng, S., Lee, K.T., So, Y.S., Hong, J., Choi, J., Byun, H.J., Kim, H., Bang, S., et al. (2015). Systematic functional profiling of transcription factor networks in Cryptococcus neoformans. Nat Commun 6, 6757.

42. Botts, M.R., Giles, S.S., Gates, M.A., Kozel, T.R., and Hull, C.M. (2009). Isolation and characterization of Cryptococcus neoformans spores reveal a critical role for capsule biosynthesis genes in spore biogenesis. Eukaryot Cell 8, 595–605.

43. Heitman, J. (2006). Sexual reproduction and the evolution of microbial pathogens. Curr Biol 16, R711–725.

44. Ene, I.V., and Bennett, R.J. (2014). The cryptic sexual strategies of human fungal pathogens. Nat Rev Microbiol 12, 239–251.

45. Dyer, P.S., and O’Gorman, C.M. (2012). Sexual development and cryptic sexuality in fungi: insights from Aspergillus species. FEMS Microbiol Rev 36, 165–192.

46. Sun, S., Coelho, M.A., David-Palma, M., Priest, S., and Heitman, J. (2019). The Evolution of Sexual Reproduction and the Mating-Type Locus: Links to Pathogenesis of Cryptococcus Human Pathogenic Fungi. Annu Rev Genet.

47. Phadke, S.S., Feretzaki, M., and Heitman, J. (2013). Unisexual reproduction enhances fungal competitiveness by promoting habitat exploration via hyphal growth and sporulation. Eukaryot Cell 12, 1155–1159.

48. Ni, M., Feretzaki, M., Li, W., Floyd-Averette, A., Mieczkowski, P., Dietrich, F.S., and Heitman, J. (2013). Unisexual and heterosexual meiotic reproduction generate aneuploidy and phenotypic diversity de novo in the yeast Cryptococcus neoformans. PLoS Biol 11, e1001653.

49. Coelho, M.A., Bakkeren, G., Sun, S., Hood, M.E., and Giraud, T. (2017). Fungal Sex: The Basidiomycota. Microbiol Spectr 5.

50. Hull, C.M., Davidson, R.C., and Heitman, J. (2002). Cell identity and sexual development in Cryptococcus neoformans are controlled by the mating-type-specific homeodomain protein Sxi1alpha. Genes Dev 16, 3046–3060.

51. Lengeler, K.B., Fox, D.S., Fraser, J.A., Allen, A., Forrester, K., Dietrich, F.S., and Heitman, J. (2002). Mating-type locus of Cryptococcus neoformans: a step in the evolution of sex chromosomes. Eukaryot Cell 1, 704–718.

52. Alby, K., and Bennett, R.J. (2011). Interspecies pheromone signaling promotes biofilm formation and same-sex mating in Candida albicans. Proc Natl Acad Sci U S A 108, 2510–2515.

53. Niklas, K.J., Cobb, E.D., and Kutschera, U. (2014). Did meiosis evolve before sex and the evolution of eukaryotic life cycles? Bioessays 36, 1091–1101.

54. Goodenough, U., and Heitman, J. (2014). Origins of eukaryotic sexual reproduction. Cold Spring Harb Perspect Biol 6.

55. Umen, J., and Coelho, S. (2019). Algal Sex Determination and the Evolution of Anisogamy. Annu Rev Microbiol 73, 267–291.

56. Tanaka, R., Taguchi, H., Takeo, K., Miyaji, M., and Nishimura, K. (1996). Determination of ploidy in Cryptococcus neoformans by flow cytometry. J Med Vet Mycol 34, 299–301.

57. Wang, L., Zhai, B., and Lin, X. (2012). The link between morphotype transition and virulence in Cryptococcus neoformans. PLoS Pathog 8, e1002765.

58. Fan, Y., and Lin, X. (2018). Multiple Applications of a Transient CRISPR-Cas9 Coupled with Electroporation (TRACE) System in the Cryptococcus neoformans Species Complex. Genetics 208, 1357–1372.

59. Skene, P.J., and Henikoff, S. (2015). A simple method for generating high-resolution maps of genome-wide protein binding. Elife 4, e09225.

60. Langmead, B., and Salzberg, S.L. (2012). Fast gapped-read alignment with Bowtie 2. Nat Methods 9, 357–359.

61. Li, H., Handsaker, B., Wysoker, A., Fennell, T., Ruan, J., Homer, N., Marth, G., Abecasis, G., Durbin, R., and Genome Project Data Processing, S. (2009). The Sequence Alignment/Map format and SAMtools. Bioinformatics 25, 2078–2079.

62. Zhang, Y., Liu, T., Meyer, C.A., Eeckhoute, J., Johnson, D.S., Bernstein, B.E., Nusbaum, C., Myers, R.M., Brown, M., Li, W., et al. (2008). Model-based analysis of ChIP-Seq (MACS). Genome Biol 9, R137.

