## Supplemental figures 1-5 for "Elucidation of the determinant for orchestration of solo unisexual cycle in an important human fungal pathogen"

[1]State Key Laboratory of Mycology, Institute of Microbiology, Chinese Academy of Sciences, Beijing 100101, China. [2]University of Chinese Academy of Sciences, Beijing 100049, China. [3]University of Science and Technology of China (USTC, Hefei 230026, China. [4]Public Technology Service Center, Institute of Microbiology, Chinese Academy of Sciences, Beijing 100101, China. [5]Department of Microbiology, School of Basic Medical Sciences, Peking University Health Science Center, Beijing 100191, China.

[†]These authors contributed equally to this work.
*

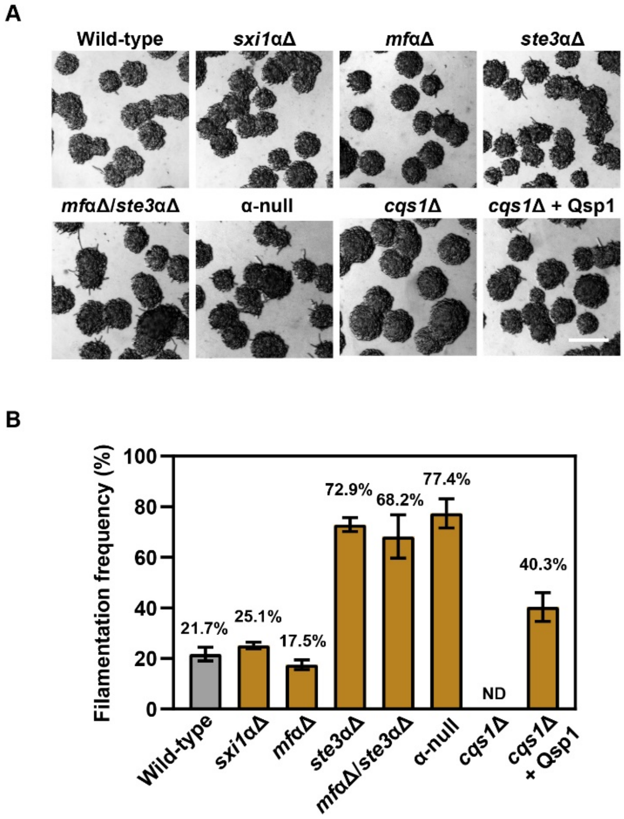


**Figure S1. α sex-determination is not essential for filamentous initiation.** The mini-colony morphology of different strains. Cells were spotted onto V8 agar at a low cell density, images were taken 21 hours after incubation (**A**). Scale bar, 100 μm. Filamentation frequency was calculated based on the percentage of filamentous mini-colonies (**B**). Data are presented as the mean ± SD from three independent experiments.

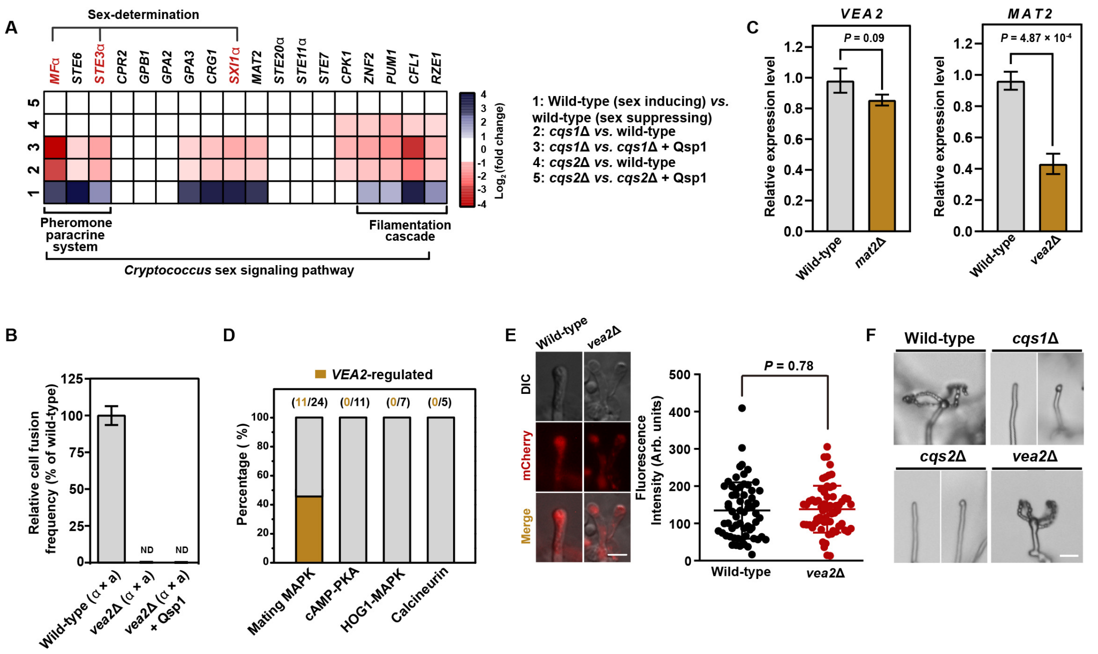


**Figure S2. Vea2 is essential for bisexual syngamy but is not strictly required for unisexual reproduction.** **(A)** Heatmap analysis indicating the expression of genes belonging to the sex signaling pathway in different *C. neoformans* strains under different conditions. The color bar represents $\log_2$ fold change. The result was based on reanalyzing published RNA-seq data [39]. **(B)** Addition of synthetic Qsp1 could not rescue the cell-cell fusion defect of *vea2*Δ mutant. The final concentration of synthetic Qsp1 is 512 nM. Data are presented as the mean ± SD of six independent experiments. **(C)** qRT-PCR shows the expression of *MAT2* and *VEA2* in *vea2*Δ and *mat2*Δ, respectively. RNAs were extracted from different strains cultured on V8 agar for 12 hours. Data are presented as the mean ± SD of three independent experiments, two-tailed Student's *t*-test. **(D)** *VEA2*-regulated genes are enriched in the mating MAPK pathway. In contrast, *VEA2*-regulated genes are not involved in the cAMP-PKA, *HOG1*-MAPK and calcineurin pathways. **(E)** The expression of Dmc1-mCherry in the basidia of the *VEA2* deletion mutant during unisexual reproduction. Images of the strains expressing Dmc1-mCherry were taken at 7 days after incubation on V8 agar. For each strain, 50 basidia were examined for the expression of Dmc1-mCherry. Two-tailed Student's *t*-test. Scale bar, 10 µm. **(F)** Sporulation phenotype of various strains. The wild-type and mutant strains were incubated on V8 agar for 7 days. Scale bar, 10 µm.

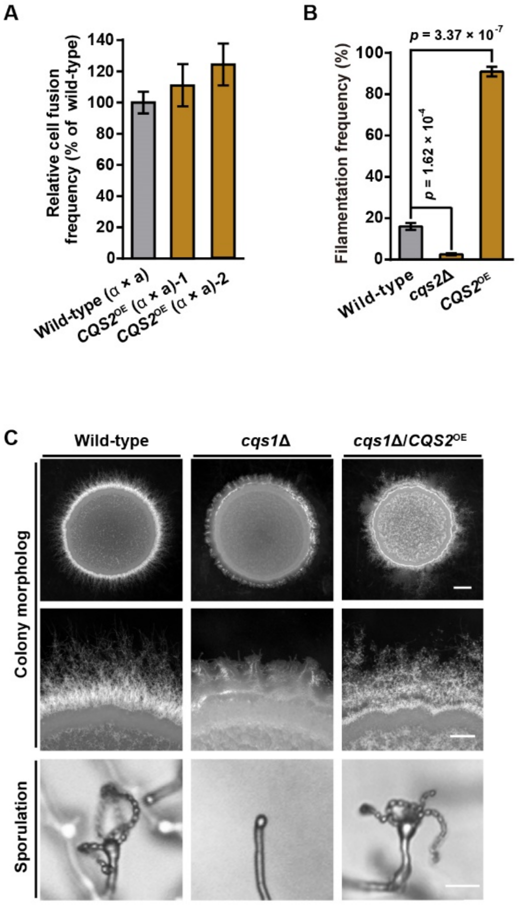


**Figure S3. Overexpression of *CQS2* restored the defects of unisexual development in the *cqs1*Δ mutant. (A)** The cell-cell fusion frequency of the *CQS2* overexpression strains compared to wild-type. The data are presented as the mean ± SD from three independent experiments. **(B)** Filamentation frequency was calculated based on the percentage of filamentous mini-colonies at 21 hours after incubation on V8 agar. The data are presented as the mean ± SD from three independent experiments. Two-tailed Student's *t*-test. **(C)** Colony morphology and sporulation phenotypes of wild-type, *cqs*1Δ and *cqs*1Δ/*CQS2*$^{OE}$. Images of different strains were taken at 7 days post inoculation on V8 agar. Scale bars, 1 mm (upper panel), 200 μm (middle panel), 20 μm (bottom panel).

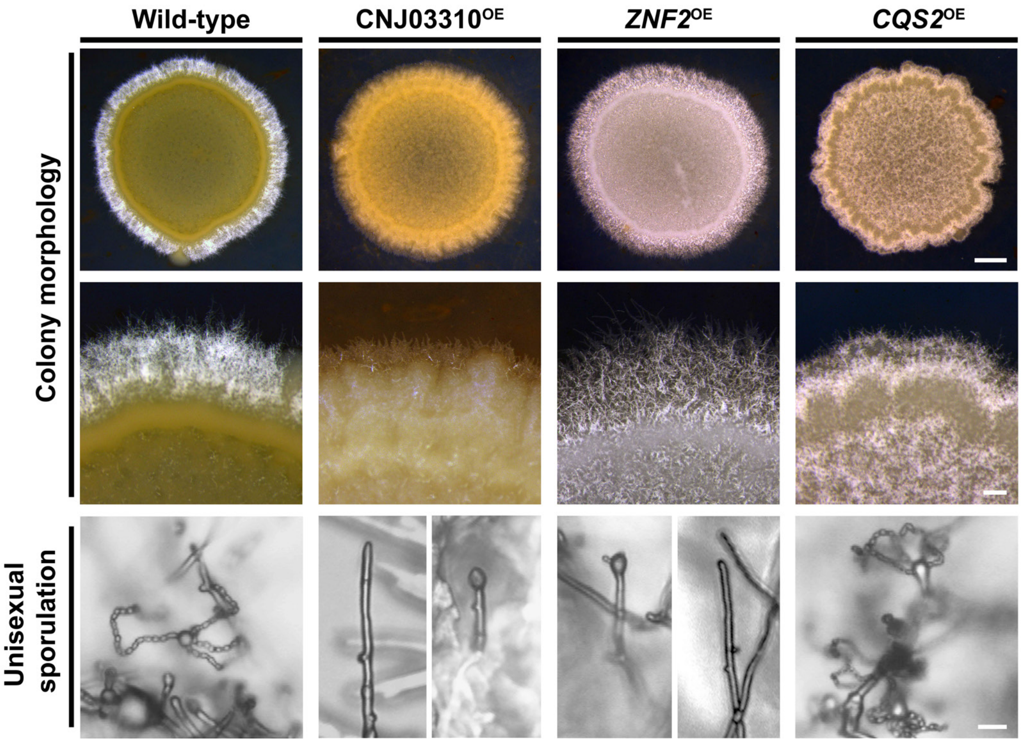


**Figure S4. Colony morphology and sporulation phenotype of wild-type, CNJ03310$^{OE}$，*ZNF2*$^{OE}$and *CQS2*$^{OE}$.** Images of different strains were taken at 7 days post inoculation on V8 agar. Scale bars, 1mm (upper panel), 200μm (middle panel), 10μm (bottom panel).

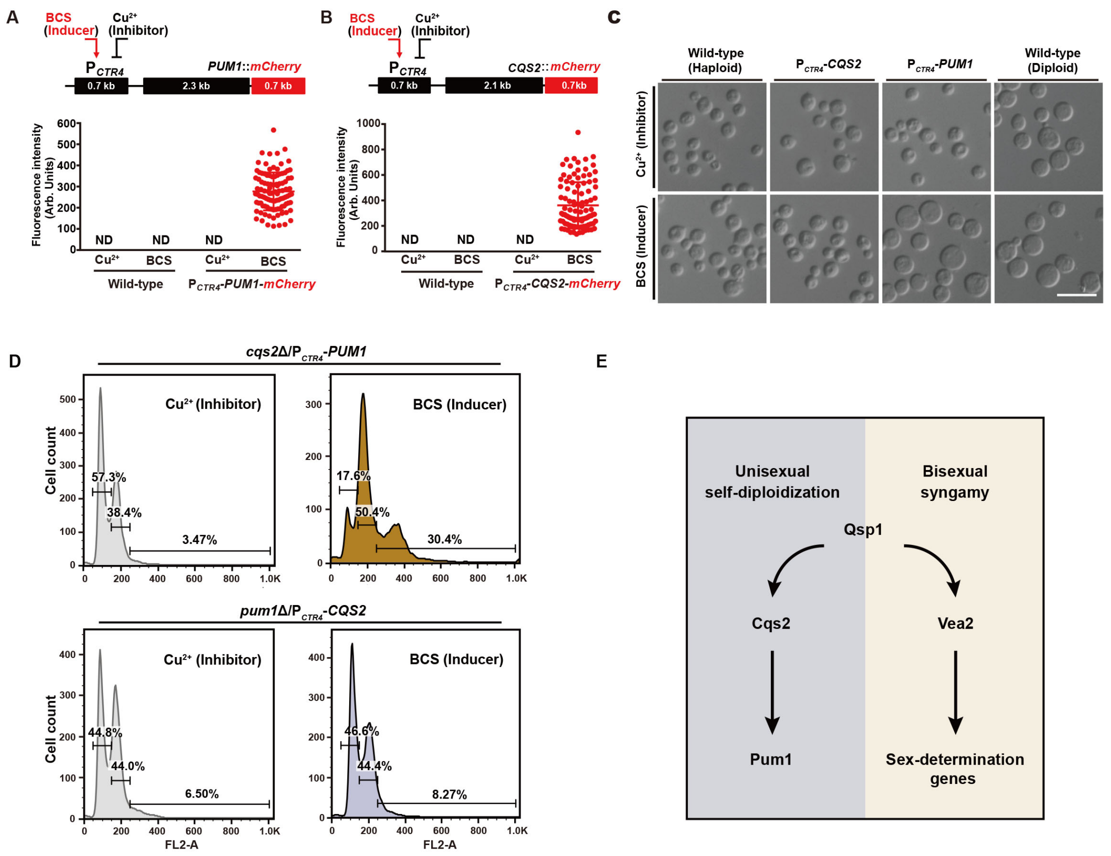


**Figure S5. Pum1 is a key target of Cqs2 during unisexual autopolyploidization. (A)** The expression of Pum1 in the $P_{CTR4}$-*PUM1*-mCherry strain under sex-suppressing condition (YPD liquid medium) in the presence of BCS or copper (n = 150 for each strain). **(B)** The expression of Cqs2 in the $P_{CTR4}$-*PUM1*-mCherry strain under sex-suppressing condition (YPD liquid medium) in the presence of BCS or copper (n = 150 for each strain). **(C)** Cellular morphology of different strains under sex-suppressing condition (YPD liquid medium) in the presence of BCS or copper. Images are representative of three independent experiments conducted with similar results. Scale bar, 10 μm. **(D)** FACS-based ploidy assessment of *cqs2*Δ/$P_{CTR4}$-*PUM1* and *pum1*Δ/$P_{CTR4}$-*CQS2* strain. Strains were cultured in YPD liquid medium for 3 days in the presence of copper (final concentration 25 μM) or BCS (final concentration 20 μM). **(E)** Model depicting that the quorum sensing peptide Qsp1 activates unisexual endoreplication and bisexual syngamy via two regulatory branches. Cqs2-Pum1 regulatory circuit drives unisexual self-diploidization and Vea2 directs bisexual syngamy via the control of sex-determination system.
